# A model of rhythm production and rhythmic auditory stimulation in healthy and Parkinsonian basal ganglia

**DOI:** 10.1101/2025.10.02.679952

**Authors:** Jacob Duda, Jonathan Cannon

**Affiliations:** Department of Psychology, Neuroscience, and Behavior, McMaster University, Hamilton, Ontario, Canada; Program in Neurosciences & Mental Health, The Hospital for Sick Children, Toronto, Ontario, Canada; Department of Physiology, The University of Toronto, Toronto, Ontario, Canada

## Abstract

In fMRI experiments, the basal ganglia is consistently activated by rhythmic action and sensorimotor synchronization to a metronome, and conditions like Parkinson’s Disease that affect basal ganglia and its dopaminergic modulation are experimentally seen to affect performance on both types of task. However, it is not clear what role this circuit or dopaminergic modulation play during rhythm production and synchronization tasks. Here, we propose that the basal ganglia may specify, maintain, and adapt the tempo with which rhythmic action (e.g. finger tapping or walking) is performed. We build a model based on previous “action selection” models of the cortico-basal-ganglia loop, altered such that cortico-basal-ganglia loops correspond not to distinct actions but to a continuum of possible action tempi. During rhythm production, an initial tempo is selected by cortical input, and rhythmic action can be automatized to continue in the absence of cortical input if tonic dopamine levels in striatum are sufficiently high. When striatal dopamine is reduced, our model reproduces two key features of dopamine deprivation in Parkinson’s disease: freezing of gait, and increased variation in produced intertap intervals during rhythmic tapping. By reanalyzing data from a recent experiment with Parkinsonian patients, we confirm the model’s prediction that increased interval variability should be largely attributable to increased tempo drift (rather than, e.g., increased timekeeper noise). This model of rhythm production is the first to invoke specific features of basal ganglia circuitry. It augments existing models of action selection in basal ganglia with the addition of continuous action parameters, and in doing so provides a starting point for further modeling of action timing and rhythm in the motor system. It offers a new model of the mechanism by which rhythmic auditory stimulation supports gait in Parkinson’s patients, and makes a new, testable prediction about sensorimotor synchronization under conditions of low tonic dopamine.

## Introduction

*“Tempo drift [*…*] was actually among the very first indicators of my Parkinson’s, before I even realized it. Because I get these student course evaluations every semester, and starting about three or four years before would be this consistent thread of comments about tempo issues, like ‘he doesn’t keep the tempo’ [*…*] and I was completely unaware of it*.*” -* Thomas Verrier, Senior Band Conductor and Director of Wind Ensembles at Vanderbilt University’s Blair School of Music, personal communication

The production of steadily paced rhythmic movement is an essential capacity that is nearly universal in the animal kingdom. Motor rhythms ranging from the steady digestive rhythm of the lobster [1] to the lamprey’s aquatic locomotion [2] are core, evolved, survival-critical features of animal behaviour. But even considering its importance to animal survival, rhythm plays an outsized role in human behaviour. Beyond our steady bipedal gait, we engage in coordinated rhythm production in the form of music and dance, and have evolved a particular capacity and tendency to synchronize the rhythms of our behaviour with that of our peers [3,4]. Our behavioural attunement to external rhythms, especially auditory rhythms, extends well beyond its social role: routine coordination of movement with a metronome can support cognitive recovery in individuals with traumatic brain injuries [5] and improve attention in children with ADHD [6], and the very presence of a metronome can draw fluent speech from stutterers [7] and produce steady, unhampered gait in Parkinson’s patients who otherwise struggle to take a single step [8].

The human penchant for synchronization relies on two simple phenomena, both available to much simpler organisms and easily modeled: entrainment and tempo modulation. Entrainment, the alignment of periodic activity to periodic forcing at a similar tempo, is observed in systems as simple as a forced pendulum, and supports synchronization within heterogeneous groups of organisms, e.g., swarms of fireflies. Tempo modulation refers to the change of the tempo of rhythm generation by changing conditions or input, and is observed in simple central pattern generators as continuous control parameters vary [9].

In humans, these two processes are coupled in such a way that we can *tempo match*: adjust our base action tempo not just to get faster or slower, but to match a specific remembered or observed tempo. Tempo matching allows us to entrain movement to a beat at a relatively constant phase across a range of tempos (an entrained simple oscillator that does not adjust its intrinsic frequency will align to its forcing at different phases depending whether the forcing is faster or slower than its intrinsic frequency). And it allows us to continue the tempo of a rhythm after the rhythm is removed (an entrained simple oscillator will revert to its intrinsic frequency in the absence of forcing). Our capacity to match tempo is most evident in tasks like synchronizing finger tapping with an auditory rhythm and then continuing the pace, but also comes into play when we spontaneously synchronize other rhythmic actions [10,11], including our walking or running gait [12], with auditory rhythms or the rhythmic movements of peers.

The capacity to modulate, match, and sustain tempo been modeled by augmenting an oscillating or interval-generating system with a tempo variable and a tempo-adjustment rule: accelerate when action is occurring late and decelerate when it is occurring early [13–15]. In some early models, tempo matching was included more directly, by adjusting action tempo to more closely match an inter-stimulus interval [16]. However, these models did not engage directly with specific neural circuits.

Where in the brain should we begin to look for a neural substrate of tempo-matching rhythm production? The basal ganglia is an appropriate starting point. fMRI experiments have consistently shown the basal ganglia to be active during rhythmic action [17–19] (as well as beat perception [17,20–22]), and basal ganglia lesions specifically impair a patient’s ability to adjust to tempo changes while synchronizing with a metronome [23]. Dopamine in the basal ganglia seems to be especially relevant. Parkinson’s disease compromises the basal ganglia’s function by depriving the striatum (its primary input structure) of dopamine; in Parkinson’s, rhythm production is often found to be more irregular (including both finger tapping [24,25] and gait [26,27]), and rhythmic action is prone to freezing (including both finger tapping [28] and gait [29]). In Parkinson’s patients prone to freezing of gait, stride cadence is more variable than in patients who do not freeze, and more variable off dopaminergic medication than on [30]. We turn therefore to the basal ganglia, its dopaminergic modulation, and its reciprocal connections with motor areas of cortex as a starting point for modeling neural mechanisms of tempo-matching rhythm production.

What is the basal ganglia doing during rhythm production? To our knowledge, the only model that incorporates basal ganglia circuitry into behavioural timing is the Striatal Beat Frequency model[31], which describes how animals can wait for a learned time interval by initiating cortical oscillations and then waiting until the striatum detects a learned coincidence of these oscillations to act. However, this model is based on observations from tasks in which single multiple-second intervals are learned over multiple exposures, and is not designed to describe tasks requiring continuous motor production of short, identical intervals that can be flexibly accelerated or decelerated.

Instead, we turn to more general models of the motor functions of the basal ganglia. The classic GPR (Gurney, Prescott & Redgrave) model and its successors propose that the basal ganglia serves to put possible courses of action in competition with each other, and ultimately causes one to be selected while the others are suppressed [32–34]. We believe that human tempo matching starts with the problem of action selection: it requires that the participant choose one course of action, tapping at a specific desired tempo, above all other potential action plans.

Here we develop a neurophysiological model of tempo-matching rhythm production in the human brain by framing it as a type of action selection, allowing us to build on previous models of the roles of basal ganglia and its dopaminergic modulation in selecting actions. We modify these models by replacing multiple discrete, selectable courses of action with a continuum of selectable action tempos represented by a continuum of overlapping neural populations.

After the selection of an action plan, the activation of that plan must be maintained and sheltered against the disruptive influence of irrelevant cortical activity. We believe that this is essentially a problem of “automaticity,” the execution of motor sequences in the absence of cognitive control and attention (a term often invoked in the context of gait). We argue below that this function, too, is largely the responsibility of cortico-basal-ganglia circuits, and is gated by dopamine levels in basal ganglia. Following the path of a previous model of periodic action generation presented by Mannella & Baldassare [35], we model automaticity as action-specific neural activity that persists due to positive feedback mediated by a cortico-basal-ganglia loop, with a critical role of dopamine in enhancing and stabilizing the feedback strength by increasing the loop gain.

Finally, to allow for tempo-matching in this tempo-maintaining circuit, we suppose that cortical representations of periodic stimuli provide excitatory input to the specific striatal populations associated with rhythmic motor programs at the matching tempo. This is essentially an “inverse model” of action, mapping an auditory pattern onto the action that generates it, and is inspired by the observation of audio-to-motor inverse models in the avian basal ganglia circuit [36]. However, these inputs compete against the influence of the dopamine-fueled excitatory loop that stabilizes action selection against noise, and as a result ongoing actions can only match a new stimulus tempo over the course of several cycles.

Simulating this model, we demonstrate that this circuit can robustly maintain an intended or measured tempo in the absence of input and in the face of neural noise, but only to the extent that its gain is up-regulated by the presence of dopamine. With reduced dopamine, and (as a result) weaker positive feedback around the loop, the model reproduces two key phenomena observed in Parkinson’s patients and associated with a failure of automaticity: because of weaker positive feedback, rhythmic action is more irregular (e.g. its tempo wavers more because the specific tempo-action plan cannot be maintained stably) and in the extreme the tempo plan ceases to be selected at all, leading to freezing of rhythmic action. It is a key prediction of this model of tempo selection that rhythmic irregularity will be observed *as a drift in underlying tempo* (as suggested by the introductory quotation drawn from a recent interview with a Parkinsonian orchestra conductor). We test this prediction by reanalyzing an existing finger tapping data set with Parkinsonian patients and age-matched controls and confirm that the increased timing variability observed in Parkinson’s can be definitively described as tempo drift, supporting the hypothesis that this model may account for neural mechanisms of Parkinsonian symptomatology.

A second area in which the model provides convergent predictions to patient data is in the role of rhythmic auditory stimulation (RAS) in stabilizing gait. It is well established that exposure to RAS (in the form of metronome or music at an appropriate tempo) can immediately ameliorate freezing and reduce temporal variability of gait in Parkinson’s patients [8,37]. When we deliver RAS to the model under conditions of low dopamine, it prevents freezing of rhythmic action and reduces its timing variability, just as in many Parksinon’s patients.

Finally, we close with a novel prediction about the effect of tonic dopamine level on motor entrainment to auditory rhythm which may be testable in future experiments.

## Results

### Modeling action selection in basal ganglia

Our model borrows much of its structure from a model of action selection in the basal ganglia by Mannella & Baldassarre (2015) [35], which in turn borrows its core structure from the foundational GPR model by Gurney et al. (2001) [33]. Our main innovation is the gradient of tempo-specific populations, as we shall discuss below. Like Mannella & Baldassarre (2015) [35], our model consists of a cortical input (Inp), cell populations representing a cortical motor region (Ctx) and a thalamic relay (Tha), and a population corresponding to each of the following sections of basal ganglia: striatal spiny neurons with D1-type dopamine receptors (StrD1), striatal spiny neurons with D2-type dopamine receptors (StrD2), subthalamic nucleus (STN), internal and external globus pallidus (GPi and GPe). Connectivity among these populations is constrained to the three key pathways through basal ganglia (Fig 1).

**Figure 1:**
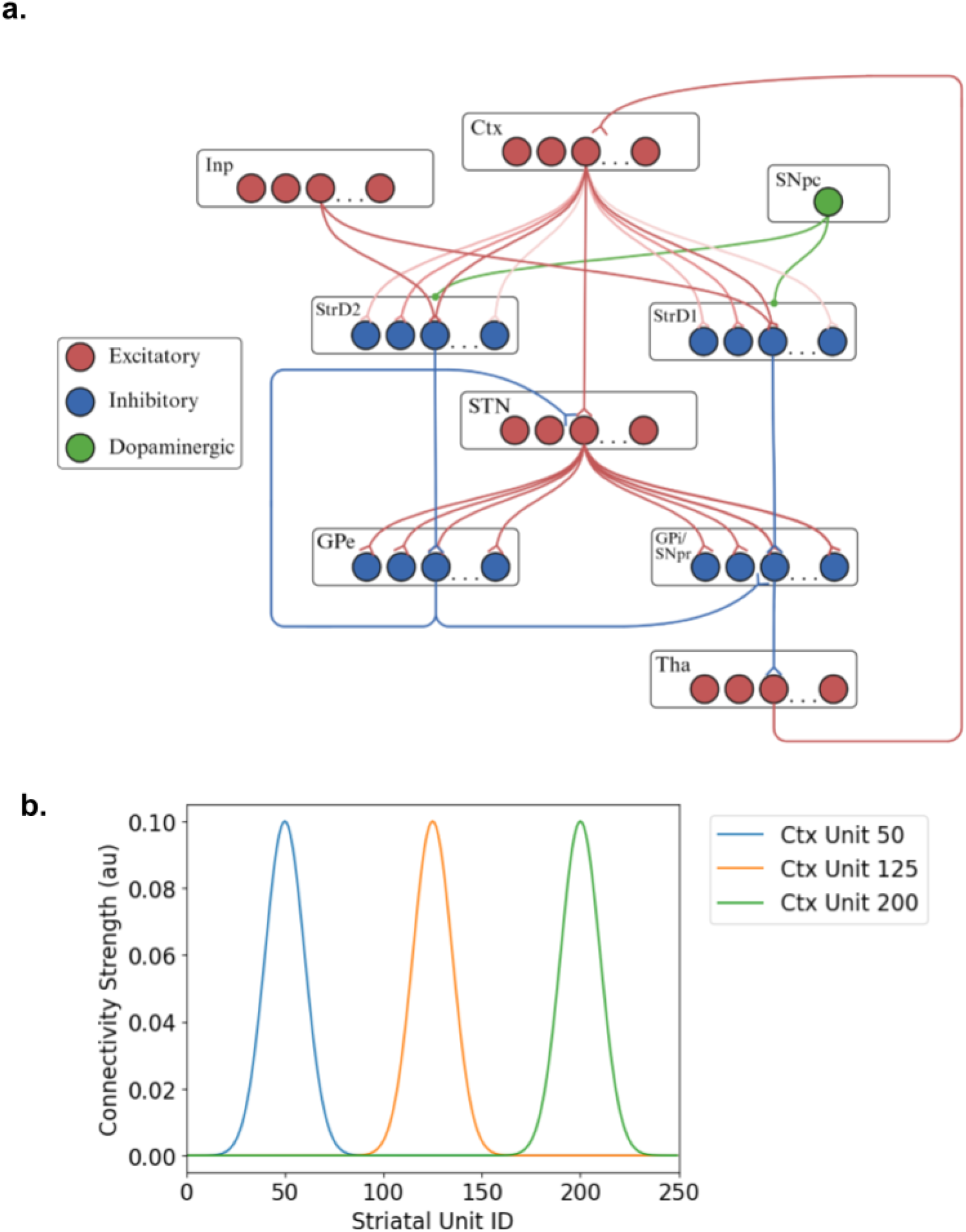
Schematic diagram of the model. a) Our model, following Mannella & Baldassarre (2015) [35], captures the proposed regional connectivity between cortex (Ctx), thalamus (Tha), and basal ganglia subregions: striatum (Str), subthalamic nucleus (STN), internal globus pallidus (GPi), and external globus pallidus (GPe), resolving these into multiple parallel sub-loops. Excitatory and inhibitory connections shown are those arising from a single sub-loop (a single unit index /preferred tempo) within the CBTC loop. Connections are segregated by sub-loop except for STN output, which is all-to-all, and connections from Ctx to Str, which are strongest within a sub-loop but extend more weakly to neighboring sub-loops as illustrated in b). This connectivity gradient leads to overlapping representations of proximal tempos in simulation.

- The *direct pathway* goes from Inp and Ctx to StrD1 (excitatory), StrD1 to GPi (inhibitory), GPi to Tha (inhibitory), and Tha to Ctx (excitatory). The double negative gives it a net excitatory (“disinhibitory”) influence on Ctx. This pathway is traditionally separated into parallel cortico-basal-ganglia-thalamo-cortical (CBTC) sub-loops, each corresponding to a possible action. Activity of a cortical input along a specific sub-loop is assumed to represent the amount of evidence that exists to suggest the associated action should be undertaken (often called “salience”). Cortical input to a sub-loop has the net effect of disinhibiting a corresponding population in thalamus, which then delivers additional excitation to the cortical population, triggering action execution.
- The *indirect pathway* adds connections from Inp and Ctx to StrD2 (excitatory), StrD2 to GPe (inhibitory), GPe to STN (inhibitory), and STN to GPi (excitatory), then continuing along the direct pathway to Tha and Ctx. The triple negative gives it a net inhibitory effect. STN projections are not segregated by action, so an increase in cortical activity encoding a specific action inhibits all possible actions similarly within the basal ganglia. STN also sends excitatory feedback to GPe.
- The *hyperdirect pathway* connects Ctx to STN (excitatory), STN to GPi and GPe (excitatory). The single negative gives it a net inhibitory effect. Due again to promiscuous projections from STN, the effect of activating this pathway is to inhibit all possible actions.

The Inp population is intended to represent input from both frontal and sensory cortical regions: the former are presumed to specify an intended motor tempo outside the context of any sensory input, whereas the latter are presumed to specify a motor tempo based on observations of time intervals periodically marked off by a rhythmic auditory stimulus, via a learned audio-motor mapping (which we do not model explicitly).

Nonspecific dopaminergic projections from substantia nigra pars compacta (SNpc) to striatum enhance the responsiveness of StrD1 neurons and reduce the responsiveness of StrD2 neurons. Gurney et al. (2001) [33] showed that the GPR model (a similar architecture without the return pathway from thalamus to cortical origin) is well suited to placing potential actions in “winner-take-all” competition with each other to the extent that dopamine is present in striatum; reduced dopamine reduces the action-specificity of the ultimate pattern of thalamic activity.

### Modeling automaticity in basal ganglia

Developments more recent than the GPR model suggest that the “actions” encoded by the parallel sub-loops actually correspond to learned sequences of action (e.g., [38]). We assume that activation of a sub-loop in the CBTC loop corresponds to an entire cyclical action sequence like a periodic finger tap or gait cycle, as discussed by Grillner et al. [39].

Like Mannella and Baldassare (2015) [35], we assume a return pathway from thalamus to the cortical population that retains segregation by action representation. This allows a particular cyclic action representation to “lock in”, in other words, neural activity corresponding to that action is initialized by action-specific cortical input but persists even in the absence of input, thus acting as an attractor in a dynamical system. Locking in an action cycle is contingent on sufficient tonic dopamine, which raises the gain around the CBTC loop, creating sufficient positive feedback for persistent activity. We hypothesize that this locking phenomenon is a mechanism for “automaticity” of the represented periodic action, defined as the continuation of the action without attention directed to the details of movement [40], and that the loss of the automatized attractor state is the dynamic that underlies freezing [29,40,41]. This hypothesis is supported by literature, especially on automaticity of human walking gait:

- Automatic performance of action is associated with reduced activity in executive and attentional cortical regions (i.e. reduced cortical input to basal ganglia) and increased functional connectivity among motor regions, including nodes in the CBTC loop [40].
- Dopamine directly modulates both freezing and automaticity: dopamine depletion/denervation reduces measures of automaticity of gait (e.g., increases stride variability and dual tasking interference) [42,43], and incidence of freezing is reduced by dopaminergic medication [44].
- Measures of reduced gait automaticity correlate with incidence of freezing of gait [45,46], suggesting a common mechanism.

### Modeling a continuum of tempo-specific CBTC sub-loops

Several lines of evidence have led researchers to suggest that the basal ganglia may be encoding “vigor,” and in particular the speed, with which an action is performed [47–50]. Our model builds on this conclusion by positing that overlapping subpopulations within the CBTC loop select for tapping or walking tempo, another continuous parameter of movement which overlaps with the more vague concept of “vigour”. Thus, rather than modeling separate subpopulations in each region corresponding to multiple qualitatively different actions, we model a large set of parallel sub-loops, each corresponding to the same action but coding for a distinct tempo. The assumption of distinct basal ganglia representations of distinct motor tempos is supported by the observation in primate striatum (putamen) of a large number of neurons tuned to specific preferred inter-tap interval durations in a rhythmic tapping task [51].

Evidence from mice indicates that neuronal representations of different movements overlap to an extent reflecting the similarity of the two movements, resulting in continuous tuning curves of neuronal sensitivity to continuous features of movement. In particular, striatal neurons show continuous tuning curves for running speed [49], and greater similarity between natural movements was associated with greater overlap of striatal population activity [52]. In this model, we create such overlapping representations and continuous tuning curves by adding cross-talk between sub-loops representing similar tempi in the form of a spread of cortical excitatory connections to striatum (Fig 1b). These connections break the strictly parallel structure of the sub-loops and excite populations corresponding to a nearby range of tempi. We assume that the combined output of the thalamus is integrated by its cortical targets, which generate cyclic activity at a tempo determined by an activity-weighted average of the preferred tempos of the thalamic units. Thus, moving forward, the tempo encoded by the network and thus produced by movement is assumed to be an activity-weighted average of the preferred tempos of the units making up the thalamic layer (see Methods for more details). However, for tempo-flexible behaviours like gait that are more controlled by subcortical central pattern generators than by learned cortical dynamics, we could equally well have assumed that the output from GPi was selecting action tempo by activating and modulating central pattern generators [39] (e.g. via the mesencephalic locomotor region [53]) with very similar results.

A neural system that can maintain any one of a continuum of states without additional input can be considered a “neural integrator”: the influence of new input that changes the state of the system is integrated and maintained over time [54]. As such, our model bears a close relationship to the picture of basal ganglia control of movement described by Yin (2023) [50]: a transient cortical input feeds into a basal ganglia integrator, and the basal ganglia output controls a lower order behavioral variable (in this case, movement cycle speed). We leave a more complete exploration of the relationship between these conceptual models to future work.

### Model behavior in simulation

In a state of normal tonic dopamine (see Methods for parameter values), and when initialized with a brief cortical input on a channel corresponding to a specific fixed tempo, the model can maintain a small bump attractor at approximately that tempo that persists when the input ceases. The units at the center of the bump in Ctx, SD1, SD2, SDN, and Tha take on higher activations, and the units at the center in GPi and GPe take on lower activations (Fig 2a). Since the preferred tempos of the units active as part of the bump attractor in Tha are assumed to set the tempo for rhythm production, the maintenance of the bump corresponds to the maintenance of a specific motor tempo. Bumps can be approximately maintained at any tempo over a wide range. We posit that the maintenance of a bump attractor in the absence of input corresponds to the “automatized” activation of a cyclic action. In the presence of cortical noise, the bump’s position drifts, causing gradual drift in the produced tempo (Fig 2b).

**Figure 2:**
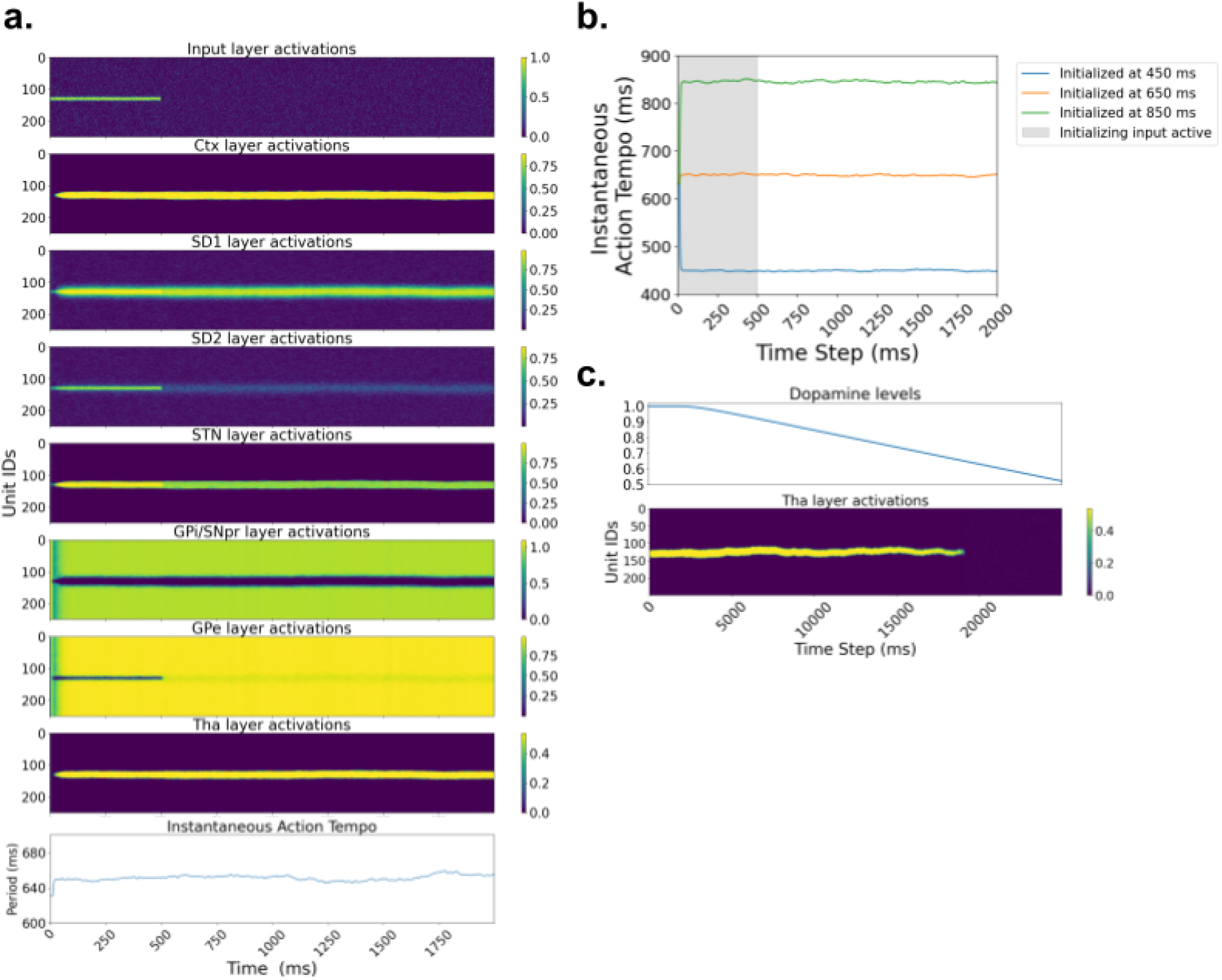
Model behaviour for various initial tempi and dopamine levels. a) Visualization of neural activity by region, plus instantaneous action tempo, over time during tempo maintenance with noise. Simulation is initialized with cortical input specifying an intended period of 650ms, corresponding to excitation centered at unit number 130 (where preferred tempo is ∼650ms). b) With sufficient amplification of recurrent excitation by dopamine, the model sustains a slowly drifting bump attractor, and can thus approximately maintain any tempo within the range of preferred tempos. Plot displays three traces of instantaneous action tempo over time after initialization with three different brief tempo-specifying inputs. c) Dopamine is necessary for continuation of rhythmic action. When dopamine level is gradually reduced in the absence of tempo-specific cortical input, the structured neural activity dies out.

By design, dopamine plays an essential role in maintaining a bump attractor by raising the gain around the CBTC loop. When the dopamine level is progressively reduced, the bump is eventually lost and the network reverts to unstructured activity (Fig 2c). We posit that this model behavior corresponds to freezing in Parkinson’s disease, which (as we discuss above) correlates with reduced measures of automaticity and occurs more often when dopamine levels are lower.

Dopamine also stabilizes instantaneous action tempo and the resulting inter-tap intervals (example simulations illustrated in Fig 3a). We conducted 100 simulations of the model at normal dopamine and 100 at a lower level dopamine (though high enough to sustain rhythmic activity), each with independent noise and lasting long enough to produce a series of 60 taps. We observed significantly higher variance in produced inter-tap interval when dopamine was low (*p* < 0.001) (Fig 3b, left).

**Figure 3:**
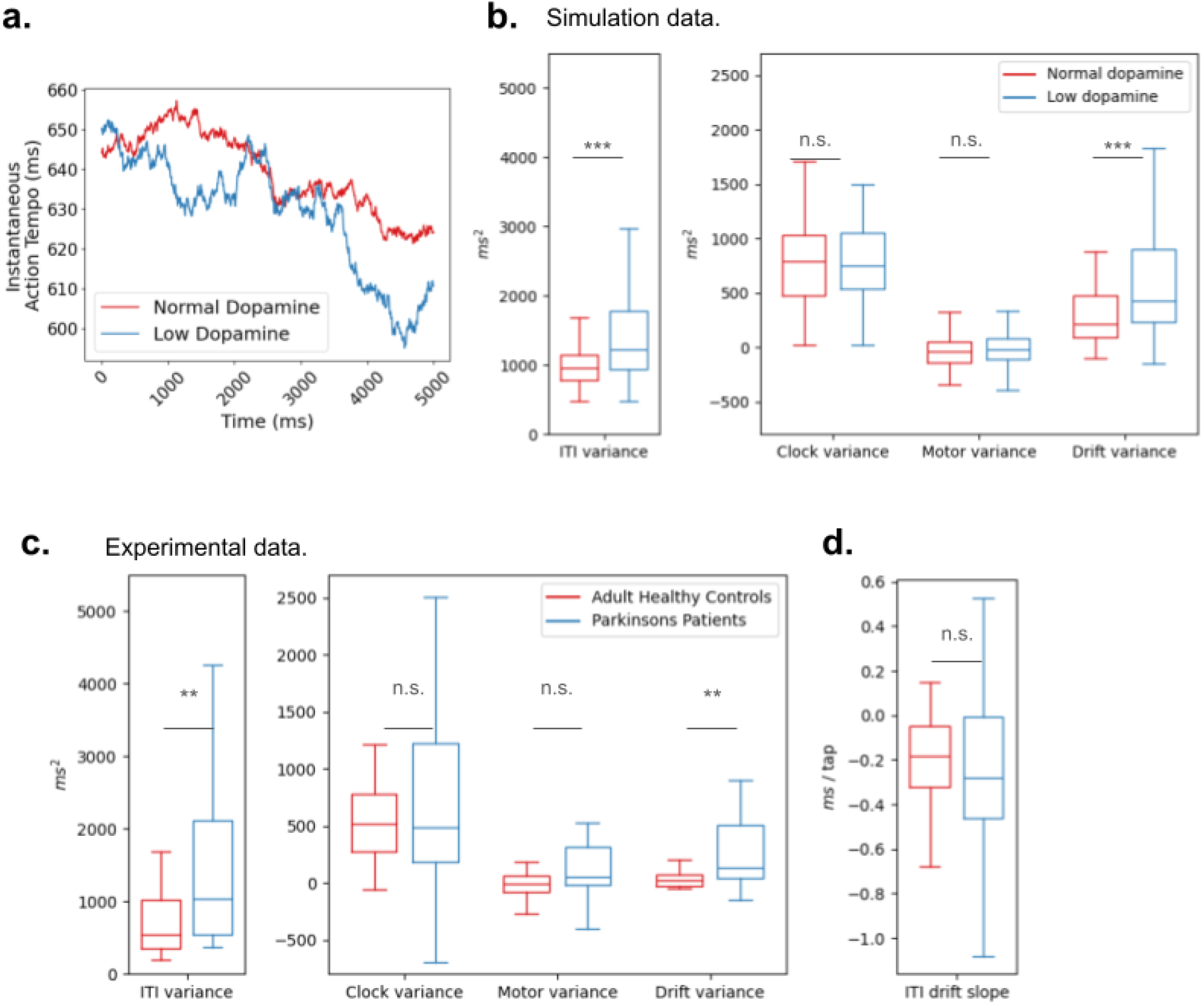
Low dopamine causes greater tempo drift in simulation and in reanalyzed human data. a) In a pair of example simulation runs with the same random seed, the network with low dopamine appears more susceptible to rapid fluctuations in the instantaneous action tempo. b) In our model, low dopamine causes an increase in inter-tap interval variance. Decomposing this variance into motor, clock, and drift variance, we find that the increase in variance can be attributed to higher tempo drift. Box plots represent 100 runs of the model in each condition with different random seeds; *** indicates p<0.001. c) In reanalyzed data comparing Parkinson’s patients to healthy controls, we find that observed differences in inter-tap interval variance are indeed attributable to tempo drift. Data is from Rose et al. (2020) [25]. Box plots represent 30 Parkinson’s patients and 26 healthy controls; ** indicates p<0.01. d) No significant difference in acceleration/deceleration trend between Parkinson’s patients and healthy controls indicates that group differences in tempo drift cannot be attributed to a tendency of Parksinon’s patients to speed up or slow down.

### Reanalysis of data from Parkinson’s patients

Elevated inter-tap interval variability was observed in Parkinson’s patients (on Levodopa medication) relative to age-matched controls in Rose et al (2020) [25], in agreement with our model (Fig 3c, left). However, there are multiple possible causes of inter-tap interval variability. According to the classic Wing & Kristofferson model of rhythm production [55], inter-tap intervals may vary due to noise in the advancement of the internal clock (“clock variance”) or due to temporally unreliable motor execution of taps (“motor variance”). Ogden & Collier (2004) [56] expand this model to allow for a third source of variance, the gradual drift in tapping tempo (“tempo drift variance”). These sources of variance can be disentangled in a given tapping sequence because they differently affect the autocorrelations of the series of inter-tap intervals (see Methods): motor variance introduces negative correlations between consecutive inter-tap intervals [55], whereas tempo drift introduces positive autocorrelations for short time lags [56].

As reported by Rose et al. (2020) [25], the variance of inter-tap intervals differed significantly between groups (*p* = 0.007), with higher variance for the Parkinson’s patients. Applying a variance decomposition to our simulation data to separate inter-tap interval variance into motor variance, clock variance, and variance due to tempo drift, we found that the only significant group difference was in tempo drift variance (Fig 3b, right); thus, the elevated variability in the low dopamine condition could be attributed to increased tempo drift. Applying the same decomposition to the data from Rose et al. (2020) [25], we found the same result: the only component of the interval variance that differed significantly between groups was tempo drift variance, which was significantly higher in the Parkinson’s patients (*p* = 0.002) (Fig 3c, center). In order to rule out the possibility that this additional drift was due to a systematic tendency of Parkinson’s patients to rush or slow down, we fit lines to the ITI sequence in each trial, averaged the slopes over each participants’ two trials, and compared the averaged slopes of the Parkinson’s patients and the age-matched controls (Fig 3d). A t-test found no significant difference between the groups (*p* = 0.86), indicating that Parkinson’s patients had no systematic tendency to get faster or slower relative to controls.

### Freezing and rhythmic auditory stimulation

Many Parkinson’s patients experience freezing during both walking [29] and other rhythmic activities like finger tapping [28]. These patients often experience substantial immediate gait benefits from RAS near their natural gait tempo: it can reduce incidence of freezing of gait [8] and reduce gait variability [37]. In model simulations run with low dopamine and noisy dopamine levels (see Methods for parameter values), we observed random freezing: the attractor bump occasionally dissipated, giving way to unstructured activity in all layers (Fig 4a). We then simulated RAS by delivering periodic input to the striatum at a period of 650ms; this input represented periodic observations of the inter-click interval, and was therefore delivered to striatal populations with preferred tempo near 650ms. A pathway by which the perception of an auditory pattern activates motor circuits that produce that pattern is a type of “inverse model,” and is justified in this system by the observation of auditory-to-motor inverse models in the basal ganglia of songbirds [36]. Simulations were performed with the same random seed, and therefore the same dynamic dopamine trace, as stimulations that previously froze. In these new simulations, freezing was not observed (Fig 4b). Finally, we compared tempo variability and inter-tap interval variability with and without RAS. A set of 100 simulations was conducted with RAS, low dopamine, and no dopamine noise (to prevent freezing). The results were compared with the inter-tap intervals from the previous simulations with low dopamine and no RAS. Figure 4c shows representative example traces of instantaneous action tempo over time: RAS generally stabilized the instantaneous action tempo close to the stimulation tempo. The interval sequences generated with RAS showed significantly less inter-tap interval variance (*p* < 0.001). This reduced variability was due entirely to reduced tempo drift (*p* < 0.001) with no significant differences in clock or motor variability (*p* > 0.05) (Fig 4d).

**Figure 4:**
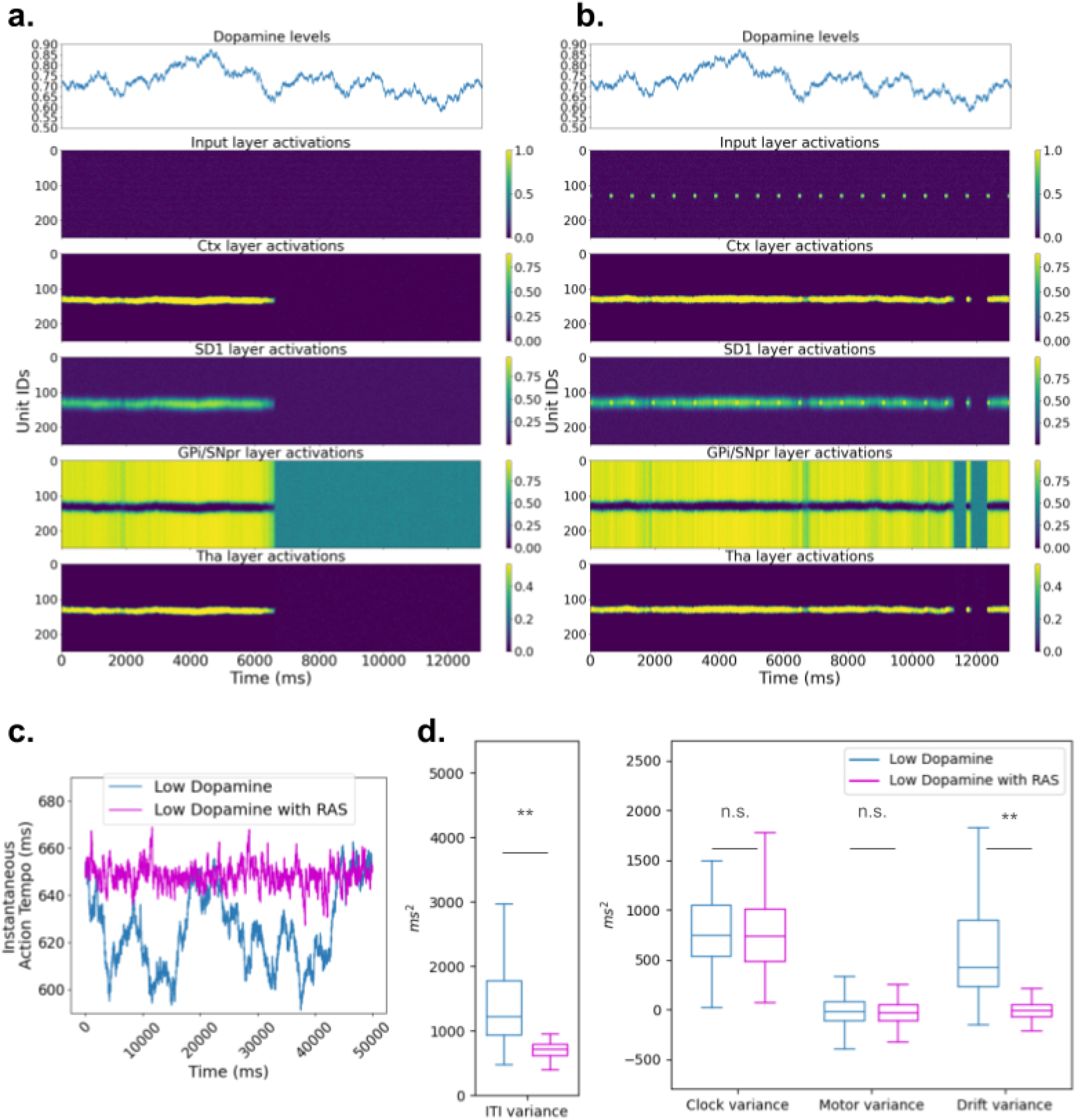
As in Parkinson’s Disease, rhythmic auditory input can prevent freezing and reduce rhythmic variability. a) When dopamine levels are allowed to fluctuate around a mean, freezing (loss of the attractor bump) is sometimes observed. b) In a simulation with the same random seed and dopamine dynamics as (a), rhythmic auditory stimulation (RAS) prevents freezing. c) RAS stabilizes tempo drift in a pair of example trials. d) Over 100 trials with and without RAS, we find that RAS reduces inter-tap interval variance, and that this effect results specifically from a reduction in tempo drift.

### Responding to tempo changes in rhythmic auditory stimulation

We next investigated the effects of dopamine level on the response to tempo changes. In a network with “normal” striatal dopamine, we delivered simulated rhythmic auditory stimulation to the network at a 650ms period (as described above) for 6 seconds and then switched the stimulation period to 700ms. The activity of the network shifted over the first next few stimulus pulses to the units representing periods in a neighborhood of 700ms (Fig 5a), correcting the tempo over the course of several taps (as is characteristic of humans tapping along with a metronome that changes tempo [57]). During the first stimulus pulse following the tempo change, the center of activation shifted smoothly from the unit with preferred tempo 650ms toward the unit with preferred tempo 700ms (Fig 5b). Next, we repeated the experiment with low dopamine. At low dopamine, we found that the network adapted more quickly to tempo changes. During the pulse immediately following the tempo change, the center of activation shifted further toward the 700ms unit when dopamine was low (Fig 5c). Similarly, at low dopamine, tapping period approached 700ms more quickly, and thus the tempo change was achieved over fewer taps (Fig 5d). Thus, our model predicts that people with Parkinson’s Disease or otherwise under conditions of low tonic dopamine will adapt more rapidly to tempo changes when trying to stay in sync with a rhythm. This prediction could be readily tested in behavioral experiments comparing conditions of lesser and greater tonic dopamine, e.g. when a Parkinsonian patient is off vs. on dopaminergic medication.

**Figure 5:**
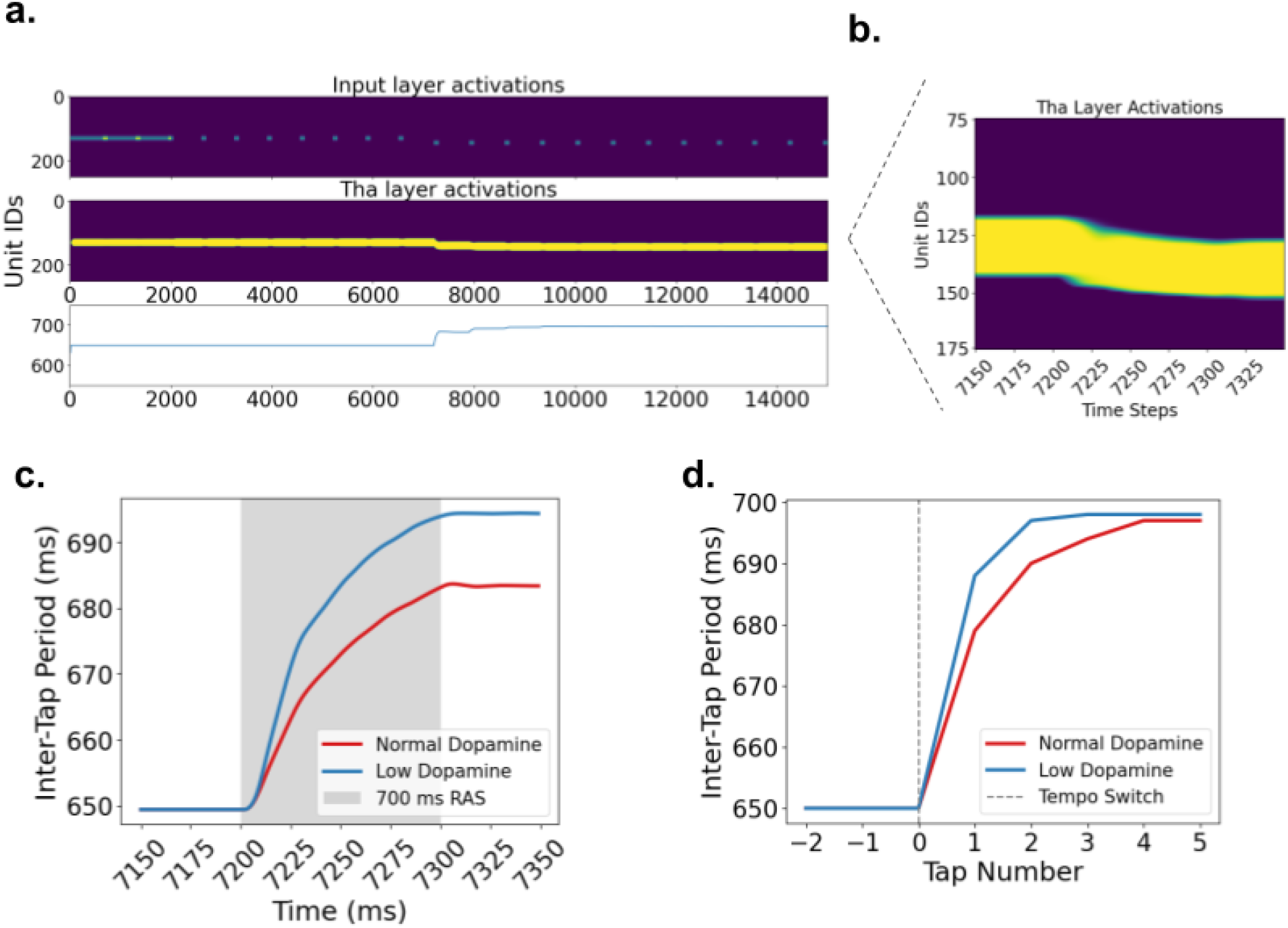
Dopamine reduction increases responsiveness to tempo changes during synchronized movement. a) The model is presented with rhythmic auditory stimulation pulses at a 650ms ISI, which then slow down to 700ms. Tempo-related cortical activity and the instantaneous action tempo determined by thalamic activity both shift to accommodate the tempo change over the next several stimulus pulses. b) Expanded view shows shifting cortical population activity over the course of a single 100ms pulse stimulus that represents the first observation of a 700ms stimulus interval. c) When dopamine level was reduced, instantaneous action tempo shifted further during the first observation of a 700ms stimulus interval. d) As a result of increased cortical responsiveness to tempo changes with reduced dopamine, the model’s inter-tap interval fully corrected to 700ms over the course of fewer taps when dopamine was lower. This behavioral prediction may be testable in simple experiments.

## Discussion

In this work, we have created a neurophysiological model illustrating a hypothesis for how a steady motor tempo is represented, automatized, and matched to a stimulus tempo in the CBTC loop. In short, overlapping subpopulations in the loop represent and actuate a continuum of tempos for rhythmic action; intentions or interval measurements from cortex act as tempo-specifying inputs to the loop; and recurrent excitation around the loop (modulated by dopamine) allows movement to persist in an “automatized” mode without tempo-specific cortical input.

The hypothesis of a continuum of tempo-specific representations in the basal ganglia is highly consistent with a long history of experimental literature on action representation and control in the basal ganglia (reviewed in Yin (2023) [50]) and integrates this literature with the observed activation of basal ganglia during deliberately paced rhythmic movement. To our knowledge, the only other mechanistic hypothesis specifically addressing the role of basal ganglia in setting the tempo for tempo-matching rhythmic action was proposed by Grossberg (2022) [58], who posited that the firing intensity, not the identity, of basal ganglia output neurons specifies the tempo for rhythmic movement. This hypothesis is somewhat at odds with ours, though both could contain a part of the larger picture. Our hypothesis and not Grossberg’s is consistent with striatal neurons whose response rates peak at specific action speeds in mice [49] and accounts for the preferred action tempos of individual neurons in the motor striatum of nonhuman primates [51]. However, the authors of the latter work also observed an increase in striatal beta oscillation amplitude as action tempo slowed, suggesting that some measure of activity in striatum changes in proportion to tempo. Note also that cells in the mesencephalic locomotor region (MLR) fire proportionately to gait speed [59–61] and are targets of basal ganglia output [53], suggesting that our “labeled lines” neural encoding of tempo may sit upstream from Grossberg’s “firing rate” encoding.

Several other relevant neural circuit models create flexible timing by providing a flexible level of input to a simple neural timekeeper that is either explicitly contained in basal ganglia [62] or speculated to reside in basal ganglia [13]. These circuit models are like Grossberg’s in that higher firing rates are associated with faster movement tempo, but different in that it is the magnitude of basal ganglia *input* rather than *output* that determines tempo. They require a constant and consistent level of input for rhythm continuation, and thus do not account for automatized rhythmic actions like normal gait and their disruption by Parkinson’s. Since dopamine modulates the gain of inputs to the basal ganglia, these models are consistent with the evidence that striatal dopamine level modulates the speed of an internal clock, at least in interval timing tasks [63]. This is certainly part of the picture of dopamine in timing; however, the effect of clock speed modulation on tempo-matching rhythm production, where both interval measurement and interval production rely in part on internal clocks, is not obvious, and this function of dopamine does not readily account for increased tempo drift in Parkinson’s patients.

In our model, greater tonic levels of dopamine make action tempo more resistant to perturbation by increasing the level of positive feedback around the CBTC loop. This feature of the model is closely related to the behaviour of the Mannella & Baldassarre [35] model on which it is based: in this model, higher levels of tonic dopamine can “lock in” a periodic action, making it difficult to change actions until dopamine level drops, permitting the selection of a new action. This proposed role for dopamine is consistent with this neurotransmitter’s apparent role in moderating between exploratory (unstable, changeable) and exploitative (stable, change-resistant) action [64–67]. However, dopamine-related resistance to tempo change has not been directly tested. Our model predicts that conditions that reduce dopamine levels should increase responsiveness to tempo changes during metronome-paced tapping. Such a finding would contrast with the observation that basal ganglia lesions decrease responsiveness to tempo changes [23]; however, the effects of lesions and dopamine deprivation may be different.

In our simulations, reduced dopamine also increases the variance of inter-tap intervals in a finger tapping task, consistent with the repeatedly reproduced observation of greater inter-tap interval variance in Parkinson’s patients [24]. According to our model, this increased variance occurs because reduced dopamine makes the representation of action tempo in the CBTC loop more susceptible to drift due to noise. Indeed, in a decomposition of inter-tap interval variance into motor variance, clock variance, and tempo drift, we find that reduced dopamine exclusively increases tempo drift. Motivated by this observation, we returned to existing experimental data and applied this decomposition to inter-tap interval sequences in Parkinsonian patients and healthy controls. Previous work claimed that increased interval variance in Parkinson’s was attributable to greater clock variance [68,69], but these analyses did not include the possibility of tempo drift, and would therefore have lumped tempo drift together with clock variance [56]. In our reanalysis, neither motor nor clock variance differed significantly between groups; only tempo drift was significantly higher in the Parkinson’s patients, as predicted by our model. We predict that treating other data sets with an analysis that separates tempo drift from clock noise would yield similar results. One caveat of this data set is that the patients were on dopaminergic medication when tested, so the tempo drift may have been related to the effects of medication rather than the effects of disease; however, several of the data sets that similarly found increased clock variance in Parkinson’s patients tested them off medication [70,71], indicating that this apparent clock variance (which we predict would be recategorized as tempo drift with the proper analysis) is a primary disease feature. If tempo drift is indeed a consistent feature of Parkinson’s, it could be used for early detection of Parkinson’s (as suggested by the introductory quotation) or for monitoring the progression of the disease.

Our model offers a new perspective on the effects of rhythmic auditory stimulation on rhythmic action. We assume that measurements of inter-stimulus intervals in a rhythmic sequence provide tempo-specific input to the CBTC loop, promoting the activation of neurons with that preferred tempo. This perspective, combined with the effects of dopamine in the model, provides a possible teleological explanation for the observation of higher dopamine levels in PET scans during tapping without a pacing signal relative to tapping with a metronome [72]: during unpaced tapping, participants must maintain a tempo without the help of additional tempo-reinforcing input from the metronome, and must therefore muster higher levels of tonic dopamine to maintain the representation of that action tempo and stabilize it against drift.

Our model also offers a new perspective on freezing of gait in Parkinson’s. In the model, freezing occurs when fluctuations in an already reduced dopamine level lead to the loss of the bump attractor state corresponding to automatized movement (generally gait) at a particular tempo. Rhythmic auditory stimulation provides repeated input specifying and reinforcing a particular tempo for action, and thus allows rhythmic action (gait) to persist through dopamine fluctuations.

Lastly, our work is to our knowledge the first modeling effort to engage with the concept of overlapping representations of action in the striatum and its role in behavior. Many movements or movement patterns are learned in the form of “generalized motor programs,” motor programs with one or more flexible parameters (for example, throwing distance or lifting force) [73]. Although we have centered our discussion around rhythmic movement, we believe many continuous parameters of movement besides periodicity may be represented in the same way. As seen in our model, a continuous representation in the input layer may allow for motor output to be adjusted smoothly in response to variable input during action plan switching or adjustment of movement (Fig 3a, Fig 5b).

### Limitations

Our model dramatically reduces the complexity of basal ganglia circuitry. In particular, it leaves out local interneurons and does not include the inputs to or dynamics of the dopamine neurons that determine both tonic dopamine level (which we specify as an input to the model) and phasic dopamine fluctuations (which are entirely left out). We do not attempt to model oscillations in the basal ganglia, which are thought to be an important contributor to Parkinson’s symptoms [74], and by using firing rate models we eliminate a wide range of possible spiking dynamics from the activity in the CBTC loop.

Importantly, we have left out the generation of neural activity cycles that presumably support cyclic action. We made this choice in order to focus exclusively on the representation of tempo. We assume that the subpopulations representing action tempo modulate the tempo of the cortical or spinal circuits that generate cyclic activity, e.g. by learned neural network mechanisms [75], central pattern generator mechanisms [76], or a combination [77]. Bringing the generation of periodicity into the scope of the model would offer a more complete picture of the neurophysiology of tempo-matching rhythmic action; this direction will be pursued in future work.

Behaviorally, our model does not provide a full account of entrainment to RAS. Classic models describe motor entrainment as supported by a combination of tempo correction, which we model here, and phase correction, which we do not. Phase correction could be added to future incarnations of the model, though accounting for its neurophysiological substrates would likely require a model of cerebellum, a structure widely implicated in phase correction [78,79]. Our model also leaves out various relevant observations on rhythm production in Parkinson’s. One is “festination,” the tendency to speed up rhythmic actions [80,81]. Parkinsonian patients (and FOG patients in particular) are especially prone to festination during slow rhythmic action [82]. Though our model does not specifically account for this phenomenon, it is compatible with festination, which could just represent the human tendency to accelerate rhythmic action, exacerbated by difficulties maintaining a constant tempo due to reduced dopamine. Alternatively, it may be related to Parkinsonian tremor [83], which is also not addressed by our model (but see [84]). Another observation not addressed here is the beneficial effects of RAS at tempos slightly exceeding the natural gait tempo [85]. This asymmetry between entrainment to faster and slower tempi may be related to asymmetries in phase and tempo correction [86–88] that we do not attempt to account for here.

## Conclusion

The model presented here provides a hypothesized account of brain mechanisms underlying tempo matching rhythm production in humans, one that we hope will guide future experimental exploration of rhythm production in the brain. Given the intimate links between motor production of rhythm, covert rhythmic time-keeping, and beat perception [89–93], we believe that our model also points toward similar hypotheses for mechanisms of beat-based rhythm perception. Future work should continue this line of thinking by continuing to engage equally with two extensive but largely disconnected literatures, one on dynamic neurophysiological modeling of the motor system and the other on behavioral and perceptual results from rhythm-related experiments in healthy and disordered human participants. We anticipate that this intersection will yield substantial progress in the quest to understand the neural dynamics underlying movement and temporal aspects of perception.

## Methods

### Mathematical formulation of model

The mathematical form of our model is borrowed from the basal ganglia circuit modeled in Mannella & Baldassarre (2015) [35]. The key differences are:

1. We do not model the cortical dynamics that generate periodic action, instead assuming that activation of the appropriate cortical population gives rise to periodic motor activity at the selected tempo.
2. The three firing rate units in each area corresponding to the three possible actions are replaced by a set of 250 units representing 250 preferred action tempi.
3. The Inp-Str and Ctx-Str connections are no longer segregated or “one-to-one”. Instead, each cortical unit is connected to the striatal layer with weights described by a Gaussian distribution (Fig 1b). Thus, if a tempo-specific input comes in from cortex, populations in the striatum with nearby preferred tempos will receive excitatory input, with populations closer in preferred tempo receiving stronger excitation.

Parameter values for the model are specified in Table 1. Parameters were tuned such that when the cortex received a strong input, corresponding thalamic units were disinhibited, and this activity could persist in the form of a line attractor (a stable “bump” of activation) when the level of dopamine is sufficiently high. Connectivity between layers was constrained to the direct and indirect basal ganglia pathways, and the sign of connections was determined by the predominant sign of the connectivity between brain regions [33,35]. Noise parameters were tuned to achieve similar levels of intertap interval variance, clock variance, and drift variance to those observed in Rose et al. (2020) [25].

**Table 1:**
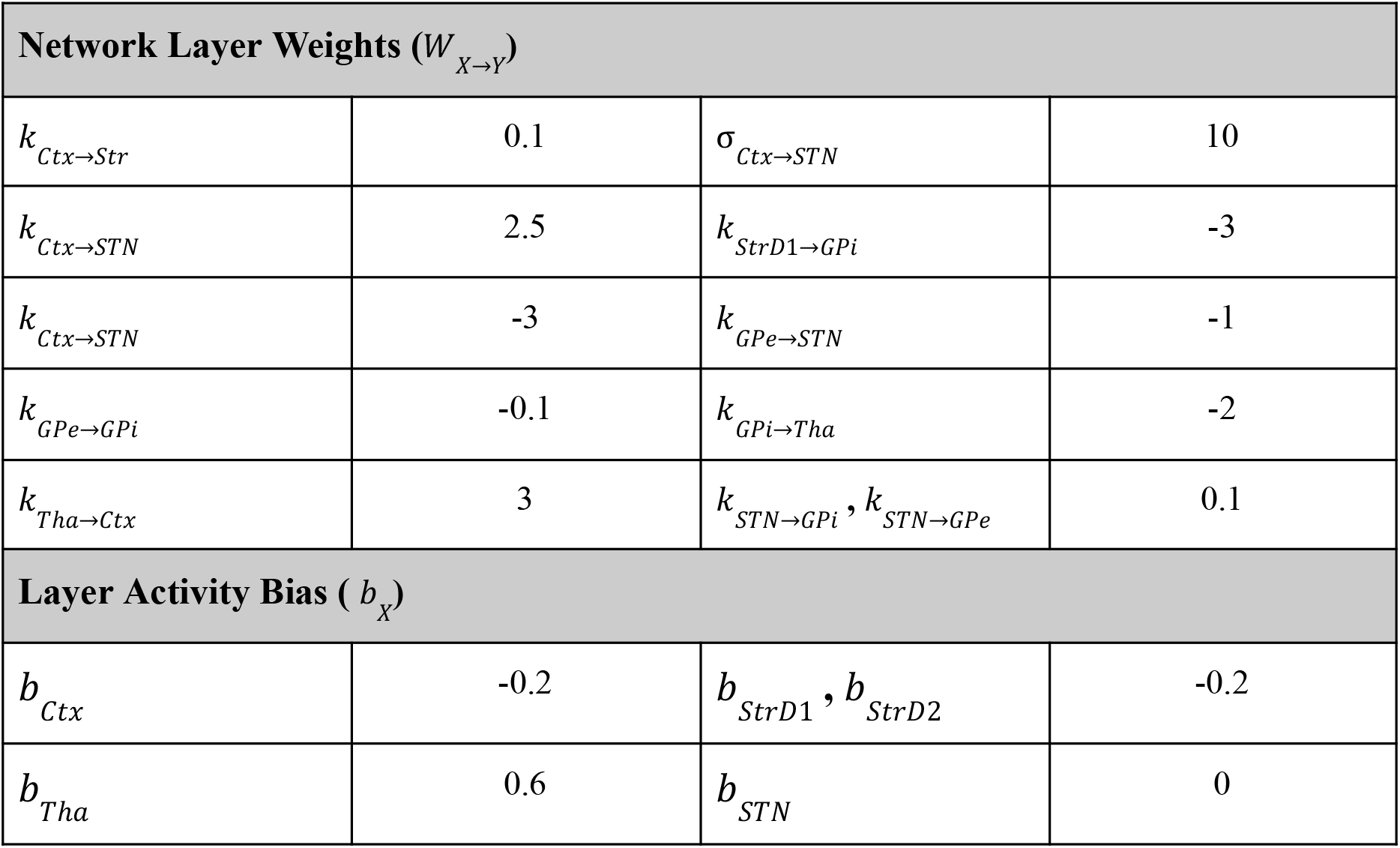
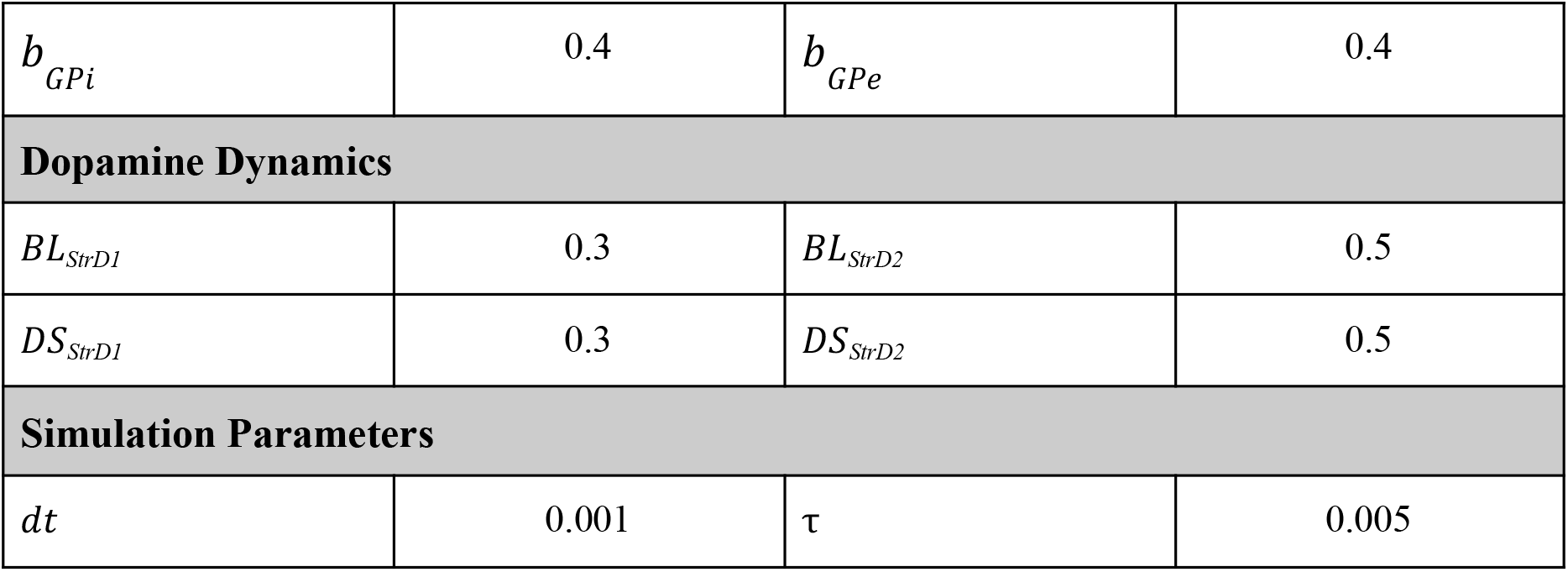
Parameters for model simulations.

Each brain area (“layer”) *X* is modeled with a vector of firing rate units whose levels of activation *A*_*X*_ follow the differential equation:

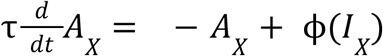

except for the Inp layer, where *A*_*Inp*_ is specified as an input to the simulation. Here *τ* is a neural time constant, *I*_*X*_ is the vector of input currents being delivered to the units at time *t*, and ϕ(*I*) = *max*(0, *tanh*(*I*)) is a nonlinear activation curve applied to each unit’s input.

Inputs to each layer follow the general equation:

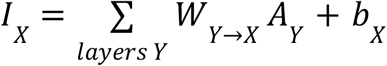

Here, *W*_*Y*→*X*_ is the matrix of weights of synapses from layer *Y* to layer *X*, which multiplies the vector of activations in layer *Y*. Bias terms *b*_*X*_ are used to tune the baseline firing rate of layers independent of external input. All weight matrices *W*_*Y*→*X*_ are set to identity matrices scaled by a constant weight factor *k*_*Y*→*X*_, representing fully segregated pathways, except for six:

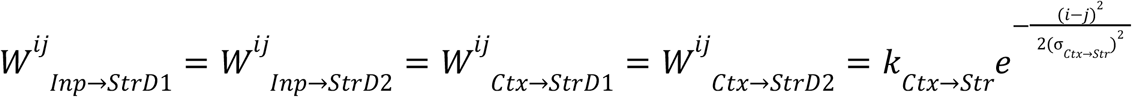

representing cross-talk between neighboring sub-loops at the cortico-striatal synapses, and

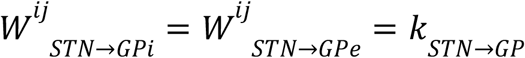

representing unsegregated, all-to-all output from *STN*.

The D1-dopamine and D2-dopamine expressing striatal layer units (StrD1 and StrD2) introduce an additional term into the expressions for *I*_*StrD*1_ and *I*_*StrD*2_ to model the effects of tonic dopamine level:

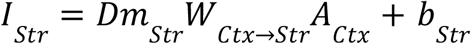

where *Dm*, the modulation of activation due to dopamine, differs between the two layers:

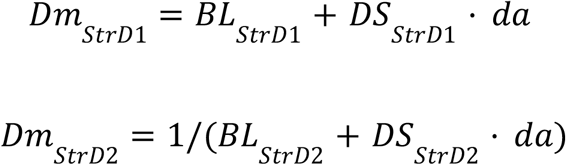

where *da* is tonic dopamine level. Thus, dopamine amplifies input gain in *StrD*1 and reduces input gain in *StrD*2. *BL* is a baseline level of modulation, while *DS* is the dopamine sensitivity of the units and controls the magnitude to which synaptic gain depends on dopamine.

The tonic dopamine level *da* evolves according to an Orstein-Uhlenbeck process with constant mean 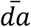:

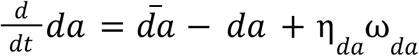

where η_*da*_ is dopamine noise level (often set to zero such that *da* rapidly approached its equilibrium 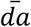) and *ω*_*da*_ was a white noise process. For a “normal dopamine” condition, 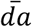 was set to 1; for a “low dopamine” condition, 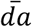 was set to 0.7.

The 250 units in each layer were assumed to represent a continuous range of tapping tempi from a 330ms period to a 1200ms period. Each unit *n* was assigned a preferred tempo *P*(*n*) (in ms) according to the formula

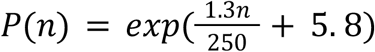

which scales more nicely than a linear or inverse map as tempo or period shrinks toward zero. The “instantaneous action tempo” that the motor system produces given a particular global activity pattern, specified by inter-tap period 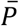 (in ms), is assumed to be determined by the “center of mass” of the unit activations in the thalamus layer:

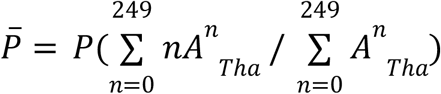

where 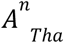 is the activation of thalamus unit *n*, such that each unit pulls the instantaneous action tempo toward its own preferred tempo proportionately to its activation.

The cortical input for each simulation was a sum of three signals: noise 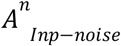, an initiation signal 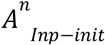, and (in some simulations) rhythmic auditory stimulation 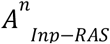. The noise source, included in all simulations except the noiseless simulations shown in Figure 5, was a white noise vector *ω* with amplitude 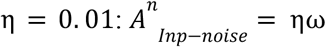. Simulations were initialized with a signal on the input units consisting of a Gaussian bump centered at the unit with preferred tempo equal to the intended initial tempo (generally 650ms, for which activation centered on unit number P^−1^): (650 *ms*) ≈ 130):

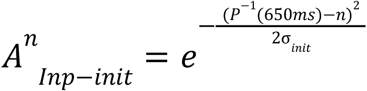

with 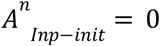 after 2 seconds (or after 0.5s in Fig 2A).

In simulations of rhythm generation under RAS, similar stimulation was delivered briefly and periodically with a 650ms period, simulating measurements of new inter-stimulus intervals:

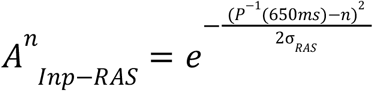

Each such stimulus input lasted 100ms, with 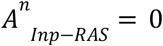 in between.

For simulations that generated tap times, a noisy countdown timer *T* was introduced that counted down to the next tap at a rate set by the instantaneous action tempo:

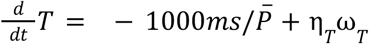

where η_*T*_ was the magnitude of timekeeper noise (introduced in order to produce “clock variability” on par with the experimental data, as discussed in Results) and *ω*_*T*_ was a white noise process as described above. Whenever *T* crossed zero, a tap was produced and *T* was reset to 1.

All differential equations were integrated with Euler’s method with *dt* = 1*ms*.

### Inter-tap interval sequence analysis

We reanalyzed a finger-tapping data in a data set collected for Rose et al (2020) [25], which found greater inter-tap interval variance during tapping continuation in PD patients relative to age-matched controls. In this data set, 30 patients and 26 age-matched controls were asked to tap their finger at a “comfortable, natural rate” with whichever hand felt more comfortable for 30 seconds in each of two trials.

For each trial’s tap sequence, we calculated inter-tap intervals and then applied the decomposition of inter-tap interval variance developed by Collier & Ogden (2004) [56] to calculate estimates of motor variance 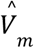, clock variance 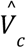, and drift variance 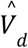:

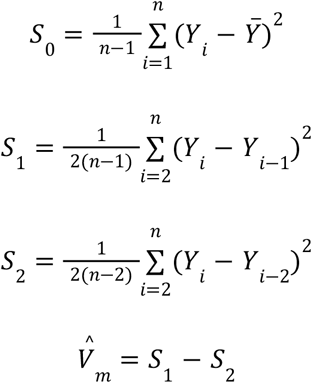

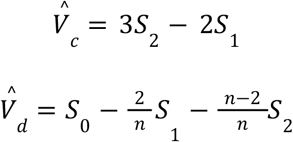

Each of these estimates was averaged over the two trials for each participant and then compared across the two groups. Since distributions of these estimates were non-normal, we compared the distributions using a Mann-Whitney-Wilcoxon test. The same analysis was applied to inter-tap intervals generated by simulations with normal and low dopamine, and then with and without RAS, as described in Results.

## Funding Declaration

This research was supported by the Human Frontier Science Program (RGEC27/2025) and a Discovery Grant from the Natural Sciences and Engineering Council of Canada (RGPIN-2022-05027).

## Acknowledgements

We are grateful to Dawn Rose and her coauthors for sharing their data and analysis code and helping us navigate it, and to John Iversen for helpful comments.

## Author contributions

J.C. and J.D. developed the theory and wrote the paper. J.D. wrote the model code. J.C. wrote the code for data reanalysis.

## Competing Interests

The authors declare no competing interests.

## Data Availability

Experimental data was taken from Rose et al. (2020) and simulated data and code is available upon request from Dr. Jonathan Cannon.

## Notes

### Competing Interest Statement

The authors have declared no competing interest.

